# Electric charge controls plasmodesma conductivity

**DOI:** 10.1101/2024.04.02.587779

**Authors:** Alexander H. Howell, Anneline H. Christensen, Vincent James, Viktoriya V. Vasina, Kaare H. Jensen, James Foley, James E. Evans, Howard A. Stone, Winfried S. Peters, Michael Knoblauch

## Abstract

While plant cells are enclosed by rigid cell walls that counteract intracellular hydrostatic pressure^1^, their plasma membrane, cytosol, and endoplasmic reticulum (ER) remain connected through plasmodesmata, nanoscopic cell wall pores^2^. Plasmodesmal cell-to-cell transport occurs in the cytosolic sleeve between the plasma membrane and the ER membrane^3–5^, and is generally thought to be limited by the size of the moving particle alone^6^. Given that biological membranes carry negative electric surface charges^7–9^, this steric notion conflicts with physical theory of ion diffusion in nanometer-sized pores with charged walls^10^. Quantifying the movements of differently sized and charged fluorescent dyes in *Tradescantia* stamen hairs, we found that anionic fluorophores of up to 1 kDa traversed plasmodesmata whereas much smaller cationic ones did not. While this agrees with theoretical expectations of different size exclusion limits for cations and anions, it questions current dogma concerning plasmodesma function and also structure, as it implies positively rather than negatively charged surfaces within plasmodesmal pores. Our findings call for re-evaluations of current models of symplasmic transport, especially of charged molecules like the phytohormone auxin (indole-acetic acid) and certain amino acids.

---

In multicellular organisms, cellular activities have to be coordinated to accomplish developmentally and physiologically meaningful results^11^. In many cases, neighbouring cells communicate via direct physical connections that establish cytoplasmic continuity^12^. Plant cells are encased in comparatively rigid cell walls, but still remain linked through cell wall pores that are traversed by cytoplasmic bridges^1‒5^. These plasmodesmata often are nanoscopic, with diameters of 50 nm or less^10^. Not only the plasma membrane and cytosol are continuous through plasmodesmata, but also the endoplasmic reticulum (ER), which forms the so-called desmotubule in the centre of the plasmodesmal pore^13^. The open space between the peripheral plasma membrane and desmotubule, the cytosolic sleeve, provides the route for symplasmic cell-to-cell transport. However, the small cross-sectional area of the sleeve, which appears further restricted by structural proteins that tether the desmotubule to the plasma membrane^14,15^, creates sterical constraints on particle movement^16^. Therefore a plasmodesma’s size exclusion limit—the size of the largest molecule capable of crossing from cell to cell through the cytosolic sleeve—is considered an essential functional characteristic^17,18^.

The significance ascribed to geometrically defined size exclusion limits in the current understanding of non-targeted (*i.e.*, purely diffusive or advective) plasmodesmal current reflects the widely held belief expressed in the title of a highly influential paper: “hydrodynamic radius alone governs the mobility of molecules through plasmodesmata”^6^. In that work, plasmodesmal conductivity was assessed through cell-to-cell movements of the fluorescent dye fluorescein isothiocyanate (FITC) conjugated to various amino acids and short peptides. While the authors did not discuss possible effects of electric charges on plasmodesmal transport, they reported all but one of their conjugates to be divalent anions at pH 8 (slightly above cytosolic pH; pH_cyt_ typically ranges between ph 7.0‒7.5); the single exception was a trivalent anion^6^. In a subsequent study, 27 similar FITC-conjugates were employed that mostly were anionic (up to pentavalent) while two were considered neutral^19^. Here the authors concluded that charge as well as molecular weight affected diffusion through plasmodesmata, but that structural features of the diffusing molecules were more important^19^. This point was later confirmed for derivatives of the green fluorescent protein (GFP), which may pass through certain plasmodesmata^20,21^. A derivative carrying additional negative charges passed more easily through plasmodesmata than one of similar molecular weight but without additional charge^22^. This result, however, appeared to be due to different hydrodynamic radii of the derivatives rather than to electric charges, and therefore seemed in line with the standard interpretation^22^. Such problems caused by complex molecular structures are mostly avoided by using comparatively simple and small fluorophores such as carboxyfluorescein or HPTS^23,24^. These tracers remain within the cytoplasm because they are negatively charged at pH_cyt_ and cannot permeate the plasma membrane^23^. For instance, carboxyfluorescein, which readily diffuses through most plasmodesmata at high rates^21,25,26^ is anionic (mostly trivalent) at pH_cyt27_.

Intrigued by the complete absence of cationic tracers from the published record of plasmodesma studies (Supplementary Table 1), we tested various small (<1100 Da) fluorophores that are differently charged at pH_cyt_. Complications due to three-dimensional tissue structure can be circumvented by studying symplasmic transport in linear cell files as found in certain trichomes (outgrowths from the epidermis)^28^; we chose stamen hairs of *Tradescantia*, classical objects in the field^19,26^, as our standard system for quantitative analyses. Shifts in intracellular hydrostatic pressure (turgor) affect plasmodesmal conductivity^29^; therefore we used diffusive injection micro-pipettes (DIMPs), which we had developed to minimize turgor-induced artifacts^30^. Analysing time-courses of fluorescence intensity in neighbouring cells, we established diffusive permeabilities of plasmodesmata for each fluorophore, demonstrating that positively charged molecules failed to move at appreciable rates while negatively charged molecules of similar and larger sizes did.

## Structure of walls with plasmodesmata

*Tradescantia* stamen hairs used for experimentation were 2.5‒4 mm long and consisted of 12 to 20 serially arranged cells (Fig. 1a). Cells were constricted towards the cross-walls between them (arrows in Fig. 1a); cell diameters at these cross-walls were 23‒42 µm with a mean of 32 µm (determined on light micrographs of 4 stamen hairs, *n* = 60). In the transmission electron microscope, the cross-walls exhibited variable thickness over short distances (Fig. 1b), and numerous plasmodesmata were visible (Fig. 1b,c). Consequently, the profile of plasmodesma lengths ranged from 170 nm to 440 nm (*n* = 94), with a modal length of about 245 nm (Fig. 1d). Based on this length profile and the dependence of diffusive current through cylindrical pores (see methods for details), a profile of the relative contributions of plasmodesmata of different lengths to total diffusive current cross the cross-walls was derived, which indicated that plasmodesmata of over 300 nm length play a minor role (Fig. 1d). The effective plasmodesma length (i.e., the length that, if it would apply to all plasmodesmata, would result in the same overall current) was 250.4 nm.

**Fig. 1.**
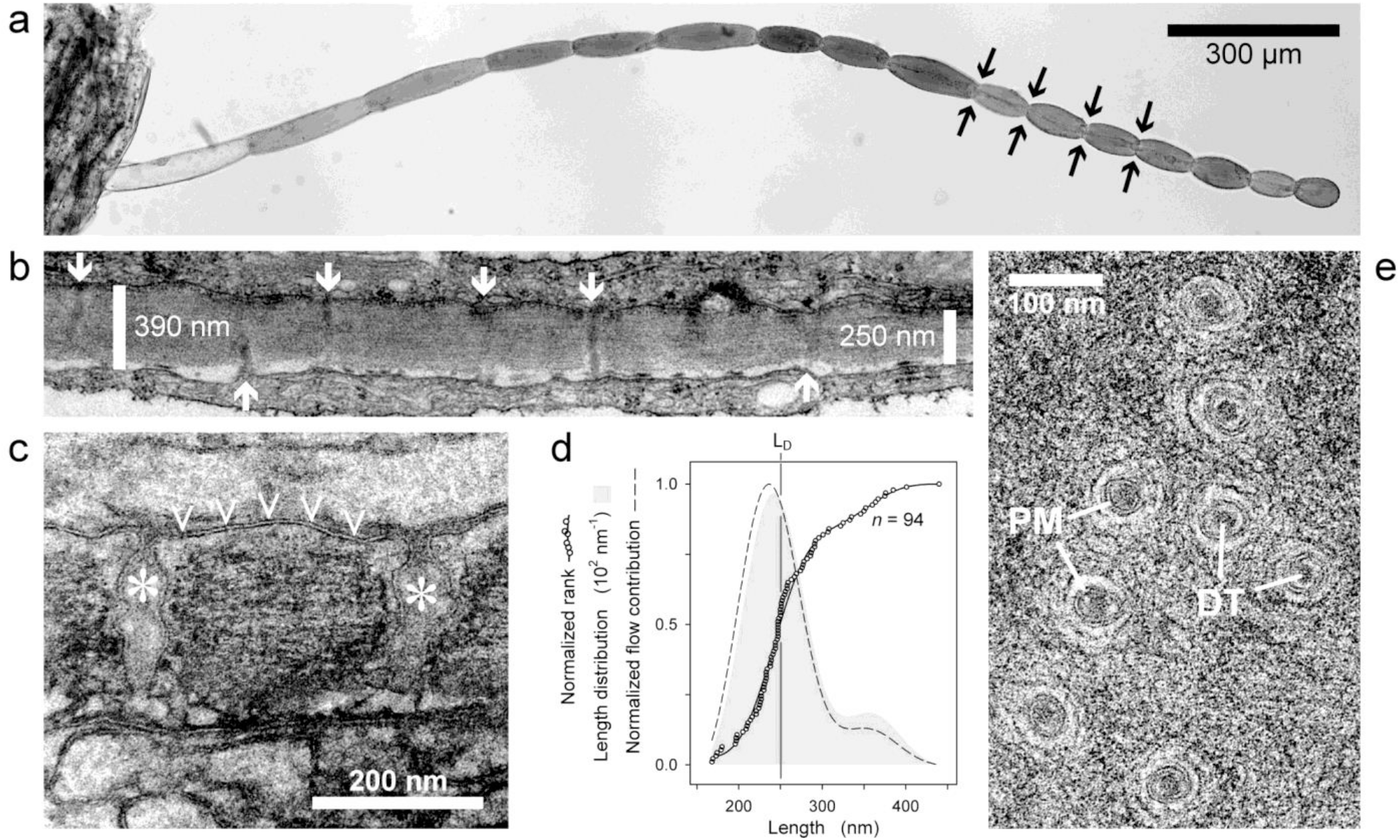
Structure of *Tradescantia* stamen hairs and the cell walls between their cells. **a**, Live stamen hair consisting of 16 serially arranged cells (light micrograph). Cross-walls between cells are located at conspicuous constrictions of the cell file (examples marked by arrows). The hair still is connected to the stamen (left). **b**, Electron micrograph of a cross-sectioned cell wall between two cells, showing numerous plasmodesmata (arrows) and variable wall thickness. **c**, Centrally sectioned plasmodesmata reveal the continuity of the plasma membrane (double line; arrowheads) into the widening and irregularly shaped plasmodesmal channels (asterisks). **d**, Dependence of plasmodesma length on wall thickness presented as the length distribution (gray shade; derived from the normalized ranks of 94 measured plasmodesmata [circles]), and the estimated relative contribution of plasmodesmata of different lengths to diffusional current between cells (dashed line). The same total current would result if all plasmodesmata had the same effective length, L_D_, of 250.4 nm. **e**, Cross-sections of plasmodesmata show the peripheral plasma membrane (PM) and central desmotubule (DT). Cell-to-cell transport occurs in the cytosolic sleeve between these two membraneous structures.

In sections more or less parallel with the plane of the transverse walls, plasmodesmata showed the plasma membrane and desmotubule, in which the ER membrane could not be identified unambiguously, and an apparently open cytosolic sleeve between them (Fig. 1e). The apparent width of the open sleeve varied significantly between about 3 and 9 nm. However, these values mostly refer to sleeve width in the interior part of the plasmodesmal pore. Close to the pore openings, pore width decreased significantly (Fig. 1c), implying much narrower sleeves in these regions. We estimated plasmodesma density to be 11.4 µm^‒2^ (*n* = 3, range 8.4 µm^‒2^‒14.4 µm^‒2^; Supplementary Fig. 1), in good agreement with an earlier study^31^. Thus, a transverse wall of average radius (16 µm; area ∼800 µm^2^) was expected to carry ∼9000 plasmodesmata.

## Fluorophore movement in *Tradescantia* stamen hairs

To visualize symplasmic transport through plasmodesmata, we introduced 19 membrane-impermeant fluorophores (Supplementary Table 2) that differed in net charge (from ‒3 to +2 at pH 7; fluorophores of net charge 0 were zwitterionic and thus membrane impermeant) and molecular mass (237 to 1027 Da) into stamen hair cells by diffusive injection, a technique that minimizes artifacts caused by artificial turgor fluctuations^30^. The hairs remained connected to the stamen during the experiments, as hair removal affects plasmodesmal conductivity^32^. Tracers frequently employed for plasmodesma studies such as Lucifer Yellow CH (charge ‒2, 443 Da) moved readily from the injected cell into adjacent ones, and so did HPTS (‒3, 455 Da), a fluorophore of superior characteristics^24^ that rarely has been used due to the lack of appropriate filters in earlier microscopes (Fig. 2). The largest anionic fluorophore tested, Alexa Fluor 633 (‒2, ∼1027 Da), remained restricted to the injected cell (Fig. 2). In contrast, none of the cationic fluorophores tested seemed able to leave the injected cell, independently of their molecular masses (237‒499 Da; Fig. 2).

**Fig. 2.**
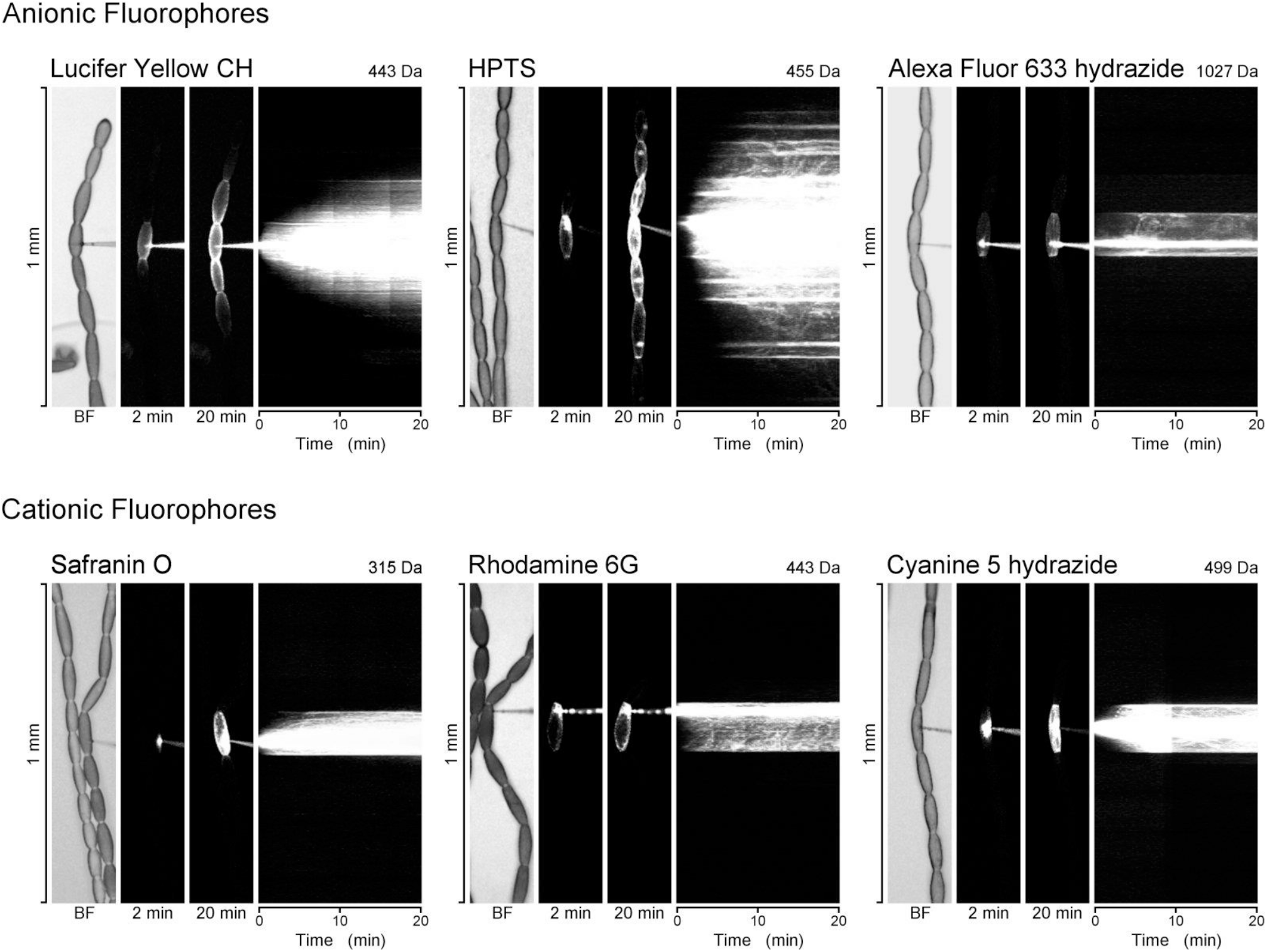
Representative diffusive injection experiments in *Tradescantia* stamen hairs. Three tests with fluorophores that are anionic at pH 7 are shown on top (molecular mass increasing from left to right) while the bottom row presents cationic fluorophores. Each panel comprises a brightfield (BF) image of the cell file at the time of impalement on the left (a diffusive-injection micro-pipette, DIMP, is impaling a cell from the right), followed by fluorescence micrographs of the cells 2 and 20 min later, and a rotated time-lapse stack (78 images) visualizing the dispersal of fluorescence over 20 min after impalement. Small anionic fluorophores diffuse along the cell file while the largest one shown (Alexa Fluor 633; ∼1027 Da) as well as cationic fluorophores in general remain restricted to the impaled cell.

To quantify the symplasmic cell-to-cell conductivity for each fluorophore, the time-courses of fluorescence intensity were determined in the injected cell and a pair of neighbouring cells on each side (Fig. 3a). Under the assumptions (i) that fluorescence intensity corresponds to fluorophore concentration, (ii) that no significant fluorophore concentration gradients exist in the cytoplasm, and (iii) that diffusive current of fluorophores occurred unidirectionally from the injected cell to the left and right, the kinetics of the fluorescence signals in three cells (*ic*‒*rn*‒*rnn* or *ic*‒*ln*‒*lnn*; Fig. 3a) allowed estimation of the symplasmic permeability of the transverse cell walls (see Methods for details). The quantitative analysis confirmed that plasmodesmal conductivity depended on electric charge while molecular masses below 1 kDa had no apparent effects (Fig. 3b, Supplementary Fig. 2). For example, the ratios of the detected permeabilities for the almost identically sized (443‒455 Da) Rhodamine 6G, Rhodamine B, Lucifer Yellow CH, and HPTS were 1: 2: 12: 43 (that is, HPTS spread from cell to cell over 40 times faster than Rhodmine 6G), corresponding to the charges of these molecules (+1, 0, ‒2, ‒3; Fig. 3b). The correlation was not perfect, though, as the monovalently anionic Sulforhodamine 101 (607 Da) did not move at notable rates, in contrast to other anionic dyes of similar sizes (Fig. 3c, Supplementary Fig. 2b). The permeability for the largest molecule tested, the divalently anionic Alexa Fluor 633 of just over 1 kDa, also was low, while the slightly smaller CF 488A (∼928 Da) moved comparatively freely. This suggested that under our experimental conditions, plasmodesmata had a size exclusion limit for anionic particles of about 1 kDa. Evidently, this limit did not apply to cationic fluorophores, as even the smallest ones tested, the cationic Acridine Yellow and Safranin O (both +2) of roughly one third the mass of CF 488A, did not travel symplasmically at noticeable rates (Fig. 3b).

**Fig. 3.**
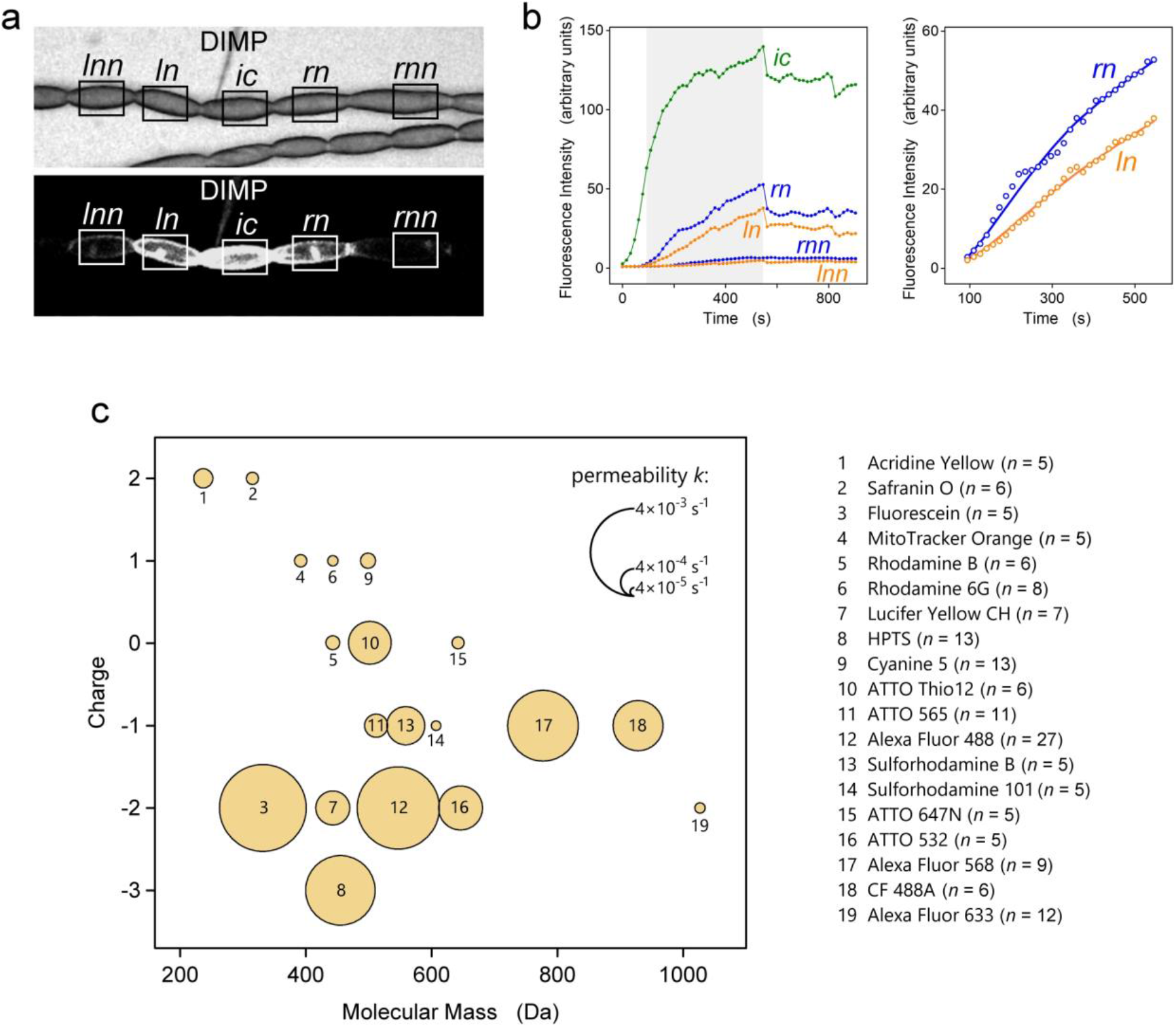
Quantification of the symplasmic permeability of transverse cell walls in *Tradescantia* stamen hairs. **a**, Live stamen hair (top, brightfield micrograph; bottom, fluorescence micrograph). A fluorophore, HTPS in this example, is injected into the cell marked *ic* by a DIMP. Its movement into the left and right neighbouring cells (*ln* and *lnn*, and *rn* and *rnn*, respectively) is monitored by quantifying fluorescence intensity in the marked rectangular fields. **b**, Time-courses of fluorescence intensity in the five cells (left graph). A model of diffusive fluorophore translocation was fitted to the data from all five cells for the period from the occurrence of fluorescence in cells *ln* and *rn* to an artificial shift in fluorescence intensity due to refocusing the microscope (grey in left graph). The model output for cells *ln* and *rn* (fitted lines on the right) provided an estimate of symplasmic permeability, *k*. **c**, The median values of the symplasmic permeabilities (*k*) determined for 19 fluorophores are represented by the areas of the circles in this plot of electric charge against molecular mass.

## Fluorophore movement in *Nicotiana* trichomes

To verify that the observed plasmodesmal selectivity for anionic compounds was not a peculiar feature of the cell type studied, we applied combinations of Alexa Fluor 488 (‒2, 547 Da) and either MitoTracker Orange (+1, 392 Da) or Cyanine 5 (+1, 499 Da) to *Nicotiana tabacum* trichome cells. The anionic fluorophore moved readily to neighbouring cells while the cationic ones remained restricted to the injected cells (Supplementary Fig. 3), in agreement with their behaviour in *Tradescantia* stamen hairs.

## Physics of electrolyte diffusion in nano-pores: implications for plasmodesmata

The significance of electrical phenomena for plasmodesmal transport has occasionally been recognized, e.g. when Tyree and Tammes in 1975 acknowledged a need to account for ‘electrostatic exclusion in the pore’^33^. However, we are not aware of experimental investigations into electrostatic effects on plasmodesma specificity and transport rates other than those discussed in the Introduction. This is surprising as a large proportion of the organic and practically all inorganic components of the cytoplasm are electrically charged, and so are the surfaces of biological membranes. The notion that the mobility of charged particles along nm-sized channels with charged walls depends on particle size alone is incompatible with physical theory and practical knowledge concerning analogous processes in artificial systems^34^. Electric surface charges within a pore containing an electrolyte solution create an electrical double layer in the solution, the thickness of which depends on the ionic strength of the solution^7^. If the thickness of the double layer is similar to or greater than the pore’s radius, electrostatic effects will control the behaviour of charged particles within the pore. A nm-sized pore with charged walls therefore may be permselective: counter-ions will be able to diffuse through the pore whereas the entrance of co-ions will be inhibited^34^. In cytosolic sleeves bordered by membrane surfaces that are negatively charged^7‒9^, one therefore should expect large conductivities for cations and significantly lower ones for similarly sized anions (Fig. 4a–d). Thus, if current notions of cytosolic sleeve dimensions and membrane surface electrostatics in plasmodesmata were correct, all of the anionic fluorescent tracers commonly used to evaluate symplasmic continuity should be excluded from plasmodesmal pores, which is contrary to what is routinely observed.

**Fig. 4.**
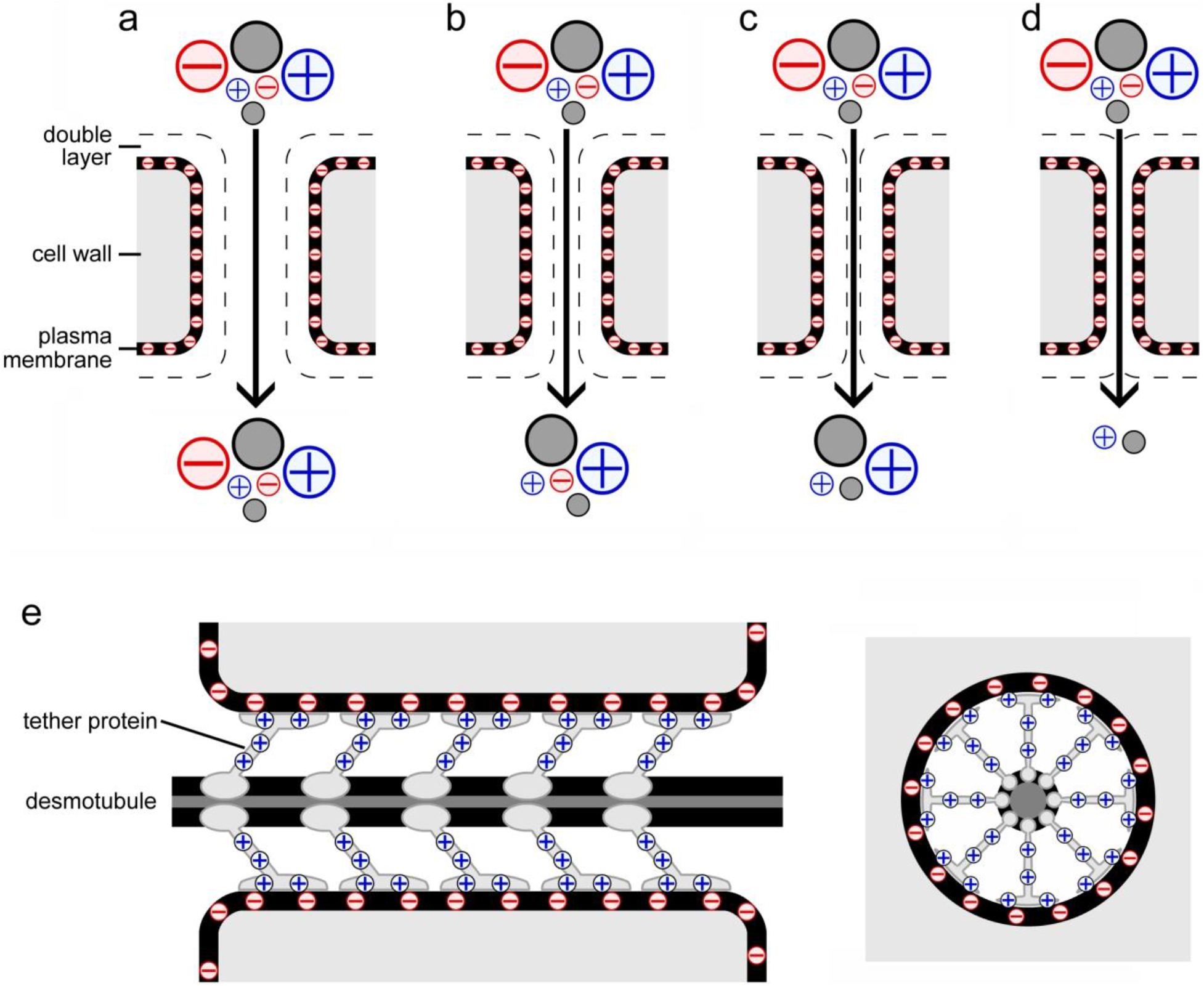
Electrostatic effects in nanoscopic pores and plasmodesmata. **a–d**, Schematic sections of cell wall pores lined with an anionic plasma membrane; pore diameters decrease from **a** to **d**. Large and small cationic, anionic, and neutral solute molecules are also shown. An electrolyte solution in contact with the negative net charge of the plasma membrane generates an electrical double layer that repels anionic solutes. This electrostatic effect will result in larger effective size exclusion limits for cations and neutral particles than for anions, which contradicts our observation of reduced plasmodesmal permeabilities for cationic fluorophores. **e**, Tether proteins known to interact with anionic membrane components may provide the positive charges along the cytosolic sleeve that would explain our results. A schematic longitudinal section (left) and a cross-section (right) of a plasmodesma with such cationic tethers are shown.

The experimental recognition and thus the awareness of electrical effects in plasmodesmata might be obfuscated by the large variability of plasmodesmal structure between cell types and developmental stages^35^. Comparatively small changes of pore size must be expected to result in qualitative modifications of the conductivities for various particles. In general, electrostatic effects of any fixed charges in the pore walls (i.e., the cytosol-exposed membrane surfaces in plasmodesmata) are negligible in pores of radii significantly wider than the electrical double layer covering the pore wall’s surface (Fig. 4a). Narrow pores in which the electrical double layers of opposing walls overlap will be penetrated preferentially by counter-ions and uncharged particles of sufficiently small size, while co-ions are excluded regardless of size (Fig. 4c,d). In intermediately sized pores, size-exclusion limits for counter-ions and co-ions may differ as they are determined sterically and electrostatically, respectively (Fig. 4b). The overall picture is further complicated by poorly understood additional effects, as suggested by the size-independent variation of the permeabilities for fluorophores of the same charge (see mono- and divalent anions in Fig. 3c). In experiments with artificial ‘ultra-nanoscale’ (<5 nm) channels, charged and uncharged organic molecules exhibit behaviors that are not predicted by current theory^36^.

Consequently, the notion that plasmodesma functionality could be quantitatively characterized by a single, geometrically defined size-exclusion limit is overly simplistic. The implications may be negligible for the behaviour of uncharged particles such as sugars flowing symplasmically to and from the phloem in source and sink organs, respectively. However, it appears questionable that symplasmic movements of, for example, anionic, cationic, and neutral amino acids proceed by the same mechanisms and with similar kinetics. Similarly, current models explaining the generation of concentration gradients across organs of charged phytohormone molecules such as auxins^37^, will have to take plasmodesmal charge selectivity into account. In live tissues, auxins like indole-3-acetic acid (IAA) are mostly conjugated to a variety of molecule types^38^, and conjugation may affect plasmodesmal permeability for auxins due to not only increased molecule size but also to changes in electric charge. Such regulation by charged conjugates of the ability to permeate plasmodesmata may occur in cytosolic molecules more generally.

## A new plasmodesma model to explain anion selectivity

The observed plasmodesma impermeability for positively charged fluorophores (Figs. 2, 3c, Supplementary Fig. 2) could be interpreted as an effect of cationic fluorophore binding to anionic membrane surfaces, which, if it occurred within plasmodesmata, could obstruct the cytosolic sleeve. Fluorophores involved in such stable associations would be helpful tools as membrane-or plasmodesma-specific fluorescent markers, but the dyes examined here lack such specificity. Moreover, when we co-injected an anionic fluorophore with a non-moving cationic one, the anionic fluorophore still was able to pass through plasmodesmata (Supplementary Fig. 3). Membrane-binding by positively charged fluorophores leading to plasmodesma blockage therefore does not explain our observations.

The experimentally observed plasmodesmal selectivity for anionic fluorophores (Figs. 2, 3c, Supplementary Fig. 2) could be explained by positively rather than negatively charged membrane surfaces within plasmodesmata. A detailed analysis based on electrokinetic theory^39^ is consistent with this interpretation (Supplementary Fig. 4). However, previous studies suggested that while the plasma membrane lipid composition in plasmodesmata differs from the norm in plant cells, the proportion of negatively charged phospholipids is similar^40^. Therefore we hypothesize that the positive charges on intra-plasmodesmal structures demanded by our results may be localized on protein rather than lipid molecules. According to current plasmodesma models, the desmotubule is tethered to the plasma membrane by proteinaceous spokes^2–4^. In general, the tethering of the ER to the plasma membrane at inter-organelle contact sites is mediated by several protein families, including proteins carrying C2 domains that bind to anionic membrane lipids through Ca^2+^-dependent or Ca^2+^-independent mechanisms^41,42^. Since the desmotubule is part of the ER, plasmodesmata represent a specialized type of plasma membrane/ER contact site^43^, and homologues of C2 proteins have been detected in plasmodesmata^44^. In contrast to regular plasma membrane/ER contact sites, the small diameters of the cell wall pores of most plasmodesmata will keep the two membranes in close proximity even without protein tethers. Such reduced functional requirements may have facilitated the evolution of new functions in plasmodesmal tethering proteins. As a working hypothesis, we suggest that the electrostatic forces responsible for the observed plasmodesmal selectivity for anionic tracers arise, first, from an at least partial shielding of the negatively charged membrane surface by the proteins, and second, from positive charges on the protein tethers that span the cytosolic sleeve (Fig. 4e). Such a scenario would open the possibility for a direct genetic control of plasmodesmal selectivity, as tether proteins with different electrical characteristics could be produced in different tissues and at different developmental stages.

It is textbook wisdom that the total concentration of inorganic cations (mostly K^+^) greatly outweighs that of inorganic anions (mostly Cl^−^) in the cytoplasm of living cells. This inorganic cation surplus balances the opposite ratio in the organic molecules, of which many are negatively charged. In this situation, positively charged pore structures favouring the entry of anions are to be expected if plasmodesma architecture evolves under selection pressure(s) that reward the free cell-to-cell movement of organic molecules. On the other hand, it often is assumed that electrical signals travel easily through symplasmic tissues^45^. However, little is known about the mechanisms of electrical signal propagation through plasmodesmata^46^, and the degree of experimentally determined electrical coupling between cells is not universally high^47^. Our results suggest that the electrical coupling between cells by the cytosolic cell-to-cell movement of relatively highly concentrated inorganic cations might be severely limited at least in some plant tissues.

## Methods

### Plants

*Tradescantia zebrina* was grown in a greenhouse at 23 °C, 60‒70% relative humidity, and a 14/10 h light/dark period (daylight with additional lamp light; 200 W full spectrum LED). *Nicotiana tabacum* var. Samsun plants were maintained in a greenhouse at 25 °C and a 16/8 h light/dark period (daylight with additional lamp light; 200 W full spectrum LED).

### Electron microscopy and analysis of plasmodesma structure

Individual stamens were excised, placed in 2% glutaraldehyde and 2% paraformaldehyde in 0.1 M cacodylate buffer, and microwave-fixed in a PELCO BioWave Pro (Ted Pella Inc., Redding CA, USA) three times for 3 min followed by 5 min at room temperature, with maximum sample temperature set to 28 °C and 750 W irradiance. After rinsing samples three times for 10 min in 0.1 M cacodylate buffer at room temperature, they were post-fixed in 1% osmium tetroxide (0.1 M cacodylate buffer) for 2 h and rinsed again three times for 10 min each. Samples were dehydrated in a methanol series (10% increments from 10% to 100%), and irradiated for 1 min in the microwave followed by 5 min at room temperature in each step. Methanol (100%) was replaced twice before transferring samples into 100% propylene oxide, which was replaced three times after 10 min each. Stamens were embedded in propylene oxide/Spurr resin at ratios of 2:1, 1:1, and 1:2 for 2 h each, and finally three times in 100% Spurr resin overnight. The samples were cured at 65 °C for 24 h. Individual stamen hairs were sectioned on a Reichert Ultracut R ultramicrotome (Leica Microsystems, Wetzlar, Germany) and sections were collected on 50 mesh PELCO copper grids (Ted Pella Inc., Redding CA, USA). For staining, sections were incubated (6 min each) in 1% uranyl acetate; ddH_2_O; 1% tannic acid; ddH_2_O; Reynolds lead; ddH_2_O. Finally, samples were imaged with a Tecnai T20 (FEI, Hillsboro OR, USA) transmission electron microscope at 200 kv.

Plasmodesma density was determined on sections of cross-walls at shallow angles (Supplementary Fig. 1). To determine plasmodesma length, longitudinal sections of stamen hairs that included cross-walls between cells were produced. A total of 94 plasmodesmata could be measured on 82 micrographs. A Savitzky-Golay spline was fitted to a plot of normalized ranks of plasmodesma lengths (Fig. 1d), the derivative of which provided a profile of plasmodesma length in the cross-walls (TableCurve2D, Systat Software, San Jose CA, USA). The diffusive current (I) of a substance through a cylindrical pore in a wall between two compartments depends on the diffusion constant of the substance (D), the concentration difference of the substance between the compartments (Δc), the cross-sectional open pore area (A), and pore length (L): I = D Δc A L^‒1^ (compare ref. ^16^). Consequently, diffusive current in pores of different lengths under otherwise identical conditions differs according to the ratios of pore lengths.

Dividing the plasmodesma length profile by L therefore provided a profile of the relative contributions of plasmodesmata of different lengths to overall current in the cross-walls, which was normalized by setting its maximum value to 1. The effective plasmodesma length, L_D_, was calculated as L_D_ = N (Σ [L_i_^‒1^])^‒1^, where N is the number of length measurements and L_i_ is the measured length of plasmodesma i.

### Confocal microscopy and microinjection experiments

Entire stamens were excised from *T. zebrina* flowers and placed on 2% w/v agar plates so that several stamen hairs lay flat on the agar surface. Agar plates were transferred to a Leica TCS SP8 confocal laser–scanning microscope (Leica Microsystems, Wetzlar, Germany) for fluorophore injection. Diffusive injection micropipettes (DIMPs) manufactured as described before^30^ were filled with one or, in most experiments, two fluorophores (0.2 mM each, in 100 mM KCl), and were mounted on an MPC200 motorized micromanipulator (Sutter Instruments, Novato CA, USA) for single-cell injection. Dye combinations were chosen from 19 fluorophores in total (Supplementary Table 2), based on their non-overlapping excitation and emission spectra. Successful injections into the cytoplasm were defined by the observation of fluorescence in the constantly moving cytoplasmic strands that traverse a cell’s central vacuole. Symplasmic movements of fluorophores were documented by time-lapse photography (minimum one image per 15.6 s) with excitation and emission settings that varied based on the fluorophores used. To visualize the dispersal of the fluorescence signals (Fig. 2), time series of fluorescence micrographs were stacked and rotated by 90° using ImageJ (https://imagej.nih.gov/ij/).

### Quantification of fluorophore movement

Total fluorescence intensities were measured in injected cells as well as in the two neighbouring cells on each side (for cell identification, see Fig. 3a) in the time-lapse image series described above (Leica LAS X, Leica Microsystems, Wetzlar, Germany). Periods for quantitative analysis were selected in which the fluorescence intensity in the impaled cell had not reached a plateau yet, and in which no artificial shifts of fluorescence intensity occurred due to cells shifting out of focus (Fig. 3b).

Assuming uniform fluorophore concentrations in the cytoplasm of each cell, unidirectional net current from the impaled cell *ic* to the left (neighbouring cells *ln*, *lnn*) and right (*rn*, *rnn*), and taking fluorescence intensity as a proxy for fluorophore concentration (*c*), the symplasmic permeability *k*_rn_ of the cell walls separating cell *rn* from its neighbours *ic* and *rnn* can be determined if time courses of fluorescence intensity, *c*(*t*), are known for the three cells.

According to a Fick’s law-like model of the concentration distribution,

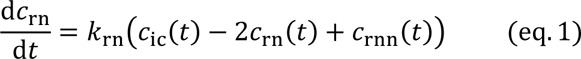

where changes of fluorophore concentration in cell *rn* over time are considered proportional to the difference in concentrations between its neighbouring cells, with the proportionality factor being *k*_rn_. The fluorophore concentration in cell *rn* is found by integrating eq. 1:

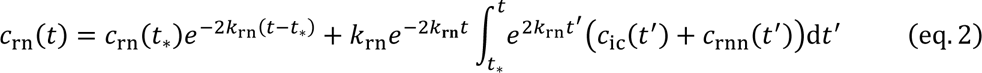

The permeability *k*_rn_ was determined by fitting eq. 2 to the observed time-course of fluorescence intensity in cell *rn*, using the experimentally determined time-courses in cells *ic* and *rnn* as input parameters, and *k*_rn_ as the only variable fitting parameter. In the process, the integral in eq. 2 was evaluated using the *trapz* function of MATLAB (v. R2020a, MathWorks Inc., Kista, Sweden). Fitting was performed using MATLAB *fminsearch* by minimizing the error function:

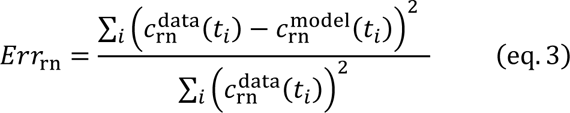

where *c*^data^ and *c*^model^ are the experimentally measured and the modelled concentration, respectively, and *t_i_* refers to the period that was analyzed. The initial estimate of *k*_rn_ required for this procedure in MATLAB was established by integrating eq. 1 and solving for *k*_rn_ with experimental concentration data from the median of:

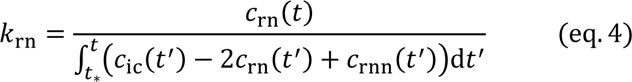

Because this method could be applied to cell *ln* analogously, we obtained two estimates (technical replicates) of the symplasmic wall permeability in every stamen hair tested, *k*_rn_ and *k*_ln_; the mean of these two values provided the *k* estimate from a given stamen hair (biological replicate). Further analyses were based on between 5 and 27 biological replicates per fluorophore.

## Data availability

Source data are provided with this paper.

## Acknowledgements

We thank Ankur Gupta and Guang Chen for discussion related to the elektrokinetic model.

## Author contributions

This project was conceived and organized by MK, HAS, and KHJ. Various aspects of the project’s experimental design and analytical procedures were developed by MK, KHJ, HAS, JEE, JF, and WSP. AHH, VJ and MK conducted injection experiments and performed electron microscopy, assisted by VVV. AHC performed the quantitative analysis of the results. WSP drafted the manuscript with input from all authors; the manuscript was discussed and finally approved by all authors.

## Additional information

### Supplementary information

The online version contains supplementary material available at https://doi.org/10.1038/―.

**Supplementary Table S1.** Small fluorescent tracers previously employed to visualize plasmodesmal permeability.

**Supplementary Table S2.** Characteristics and sources of the 19 fluorophores used in this study.

**Supplementary Fig. S1** Determination of plasmodesma density in the cell walls between stamen hair cells of *Tradescantia zebrina*.

**Supplementary Fig. S2** Dependence of plasmodesmal permeability on the molecular mass and electric charge of the 19 fluorophores tested in *Tradescantia zebrina*.

**Supplementary Fig. S3** Symplasmic movements of fluorophores injected into *Nicotiana tabacum* trichome cells.

**Supplementary Fig. S4** Comparison of experimentally observed plasmodesmal permeabilities, *k*, with theory.

