## Supplemental Tables 1-2, Supplementl Figures 1-4 for "Electric charge controls plasmodesma conductivity"

### SUPPLEMENTARY TABLES & FIGURES

**Supplementary Table 1. Small fluorescent tracers previously employed to visualize plasmodesmal permeability.** Molecular masses and electric charges of the dominant molecule form at pH 7.0 are based on currently available information and do not necessarily equal the values given in the original papers. The references represent an incomplete selection.

| Fluorophore | Mass (Da) | Net Charge (at pH 7) | Reference(s) |
|---|---|---|---|
| Fluorescein (Uranin) | 332 | −2 | 2, 10, 12, 14, 23, 25 |
| Carboxyfluorescein | 376 | −3 | 3, 4, 5, 6, 11, 13, 19, 20, 21, 24, 26, 28 |
| Lucifer Yellow CH | 443 | −2 | 2, 3, 4, 5, 8, 10, 15, 16, 18, 22, 23, 27 |
| HPTS | 455 | −3 | 28 |
| Cascade Blue hydrazide | 527 | −3 | 17 |
| Sulforhodamine G | 530 | −2 | 2 |
| Alexa Fluor 488 | 546 | −2 | 7 |
| Sulforhodamine B (Lissamine Rhodamine B) | 559 | −1 | 1, 23 |
| Sulforhodamine 101 | 607 | −1 | 18 |
| Texas Red sulfonyl chloride | 625 | 0 | 9 |
| Alexa Fluor 568 | 708 | −2 | 7 |
| Alexa 568 hydrazide | 731 | ? | 1 |
| Alexa Fluor 633 | 1150 | −2 | 7 |

10. Kragler, F. Analysis of the conductivity of plasmodesmata by microinjection. In: *Plasmodesmata: methods and protocols*, Methods in Molecular Biology 1217 (ed.: M. Heinlein). New York: Springer (2015).

**Supplementary Table 2. Fluorophores tested in this study.** Molecular masses and charges at pH 7 based on the suppliers' information (which sometimes is incomplete) and the PubChem database of the National Institutes of Health, USA (https://pubchem.ncbi.nlm.nih.gov/) are shown with the number of independent biological replicates in this study for each fluorophore.

| Fluorophore | Mass (Da) | Net Charge (at pH 7) | Supplier | Biological Replicates |
|---|---|---|---|---|
| Acridine Yellow | 237 | +2 | Thermo Scientific | 5 |
| Safranin O | 315 | +2 | Sigma Aldrich | 6 |
| Fluorescein (Uranin) | 332 | −2 | Sigma Aldrich | 5 |
| MitoTracker Orange | 392 | +1 | Invitrogen (Molecular Probes) | 5 |
| Rhodamine B | 443 | 0 | Sigma Aldrich | 6 |
| Rhodamine 6G | 443 | +1 | Sigma Aldrich | 8 |
| Lucifer Yellow CH | 443 | −2 | Sigma Aldrich | 7 |
| HPTS | 455 | −3 | Sigma Aldrich | 13 |
| Cyanine 5 hydrazide | 499 | +1 | Kerafast | 13 |
| ATTO Thio12 | 502 | 0 | ATTO-Tec | 6 |
| ATTO 565 | 512 | −1 | ATTO-Tec | 11 |
| Alexa Fluor 488 hydrazide | 547 | −2 | Invitrogen (Thermo Fisher Scientific) | 27 |
| Sulforhodamine B | 559 | −1 | Biotium | 5 |
| Sulforhodamine 101 | 607 | −1 | MedChemExpress | 5 |
| ATTO 647N | 642 | 0 | ATTO-Tec | 5 |
| ATTO 532 | 646 | −2 | ATTO-Tec | 5 |
| Alexa Fluor 568 cadaverine | 777 | −1 | Molecular Probes (Life Technologies) | 9 |
| CF 488A hydrazide | ~ 928 | −1 | Sigma Aldrich | 6 |
| Alexa Fluor 633 hydrazide | ~1027 | −2 | Invitrogen (Thermo Fisher Scientific) | 12 |

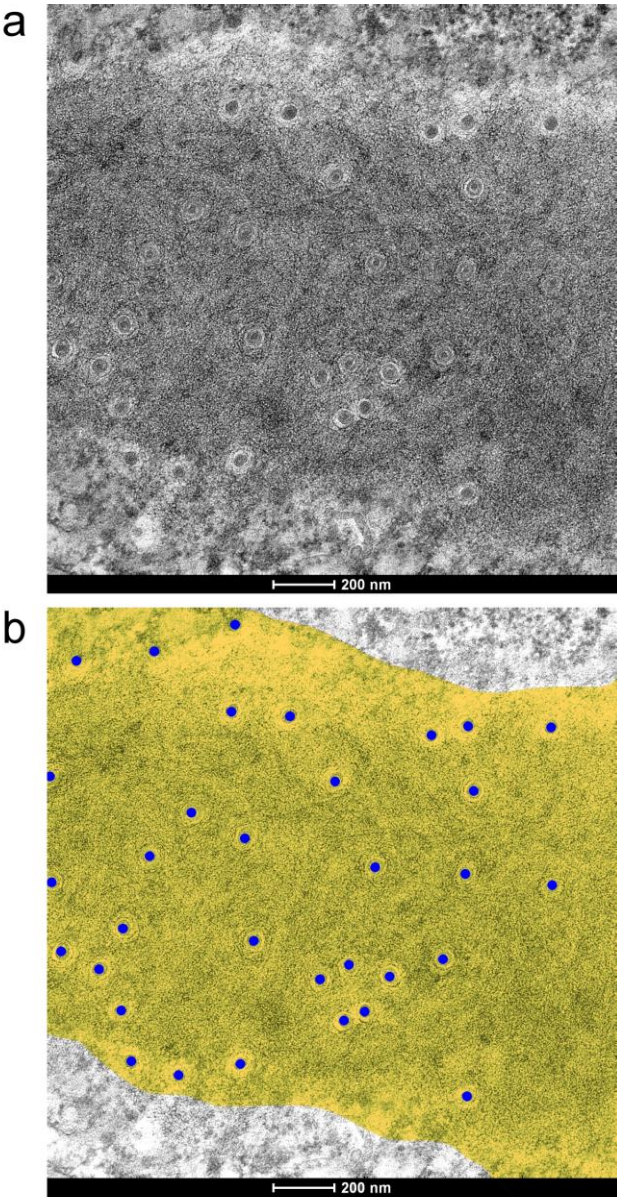


**Supplementary Figure 1. Determination of plasmodesma density in cell walls of *Tradescantia* stamen hair cells. a**, Section of a cell wall at a shallow angle showing numerous plasmodesmata (compare Fig. 1e). The total area covered by the micrograph is 3.47 $\mu m^2$. **b**, The exact location of the border between cell wall and cytoplasm is not sharply discernible on such sections and has to be estimated. The wall is marked by a yellow overlay in this image, and 33 visible plasmodesmata are highlighted by blue dots. With an estimated wall area of 2.93 $\mu m^2$, plasmodesma density is 11.3 $\mu m^{-2}$ in this image.

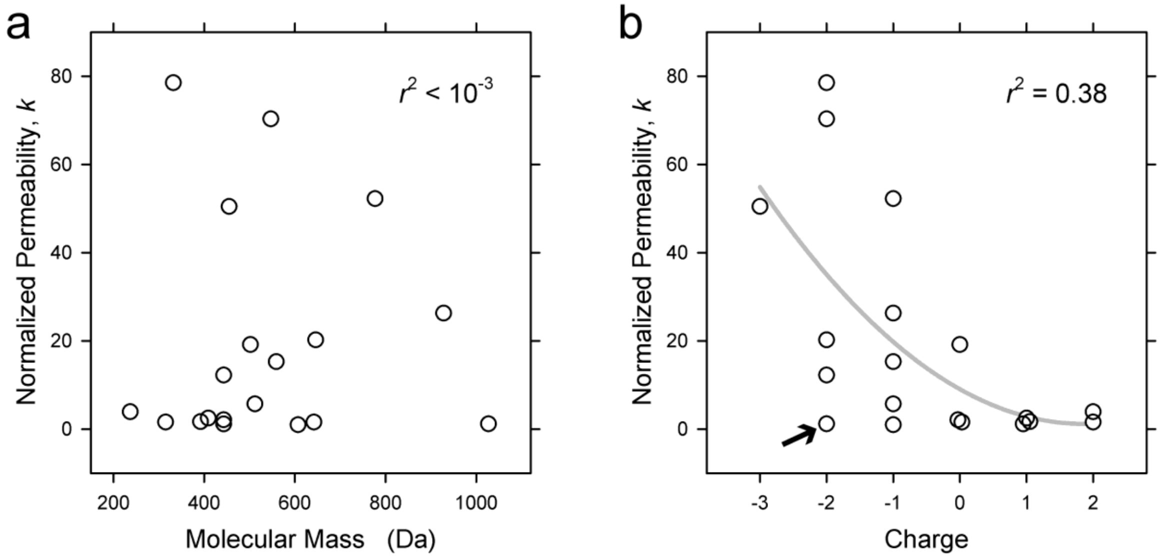


**Supplementary Figure 2. Dependence of plasmodesmal permeability *k* on molecular mass (a) and electric charge (b) of the 19 fluorophores tested.** Data shown represent the medians of the biological replicates of each fluorophore; these medians were normalized to the smallest observed value. Coefficients of determination ($r^2$) are indicated in **a** and **b**. The grey line in **b** is a second order polynomial fitted to visualize the general trend towards greater permeabilities for anionic fluorophores. The datapoint highlighted by an arrow in **b** represents Alexa Fluor 633, the largest fluorophor tested. If we assume that this large molecule is geometrically excluded and omit it from analysis, $r^2$ will rise to 0.48.

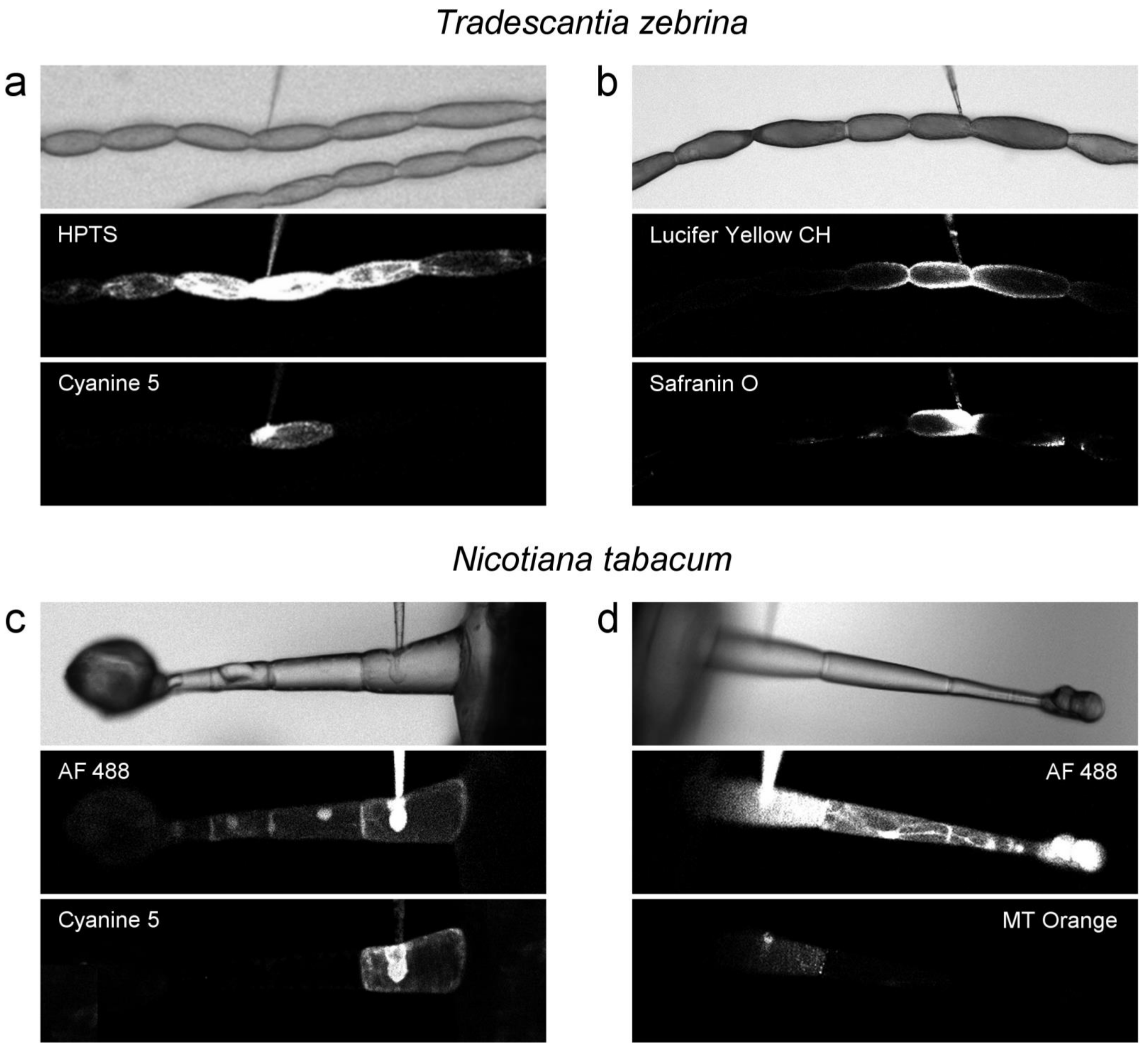


**Supplementary Figure 3. Cell-to-cell movement of anionic but not cationic fluorophores in *Tradescantia* stamen hairs and *Nicotiana* trichomes.** Combinations of one of the anionic fluorophores HPTS (**a**), Lucifer Yellow CH (**b**), and Alexa Fluor 488 (**c**, **d**) with one of the cationic fluorophores Cyanine 5 (**a**, **c**), Safranin O (**b**), and MitoTracker Orange (**d**) were injected with DIMPs into a cell in the centre of a *Tradescantia* stamen hair (**a**, **b**) or the basal cell of a *Nicotiana* epidermal trichome (**c**, **d**). After 12 minutes, all fluorophores were detectable in the injected cell, but only the anionic ones had moved into adjacent cells. Width of the micrographs: **a**, **b**, 800 µm; **c**, **d**, 450 µm.

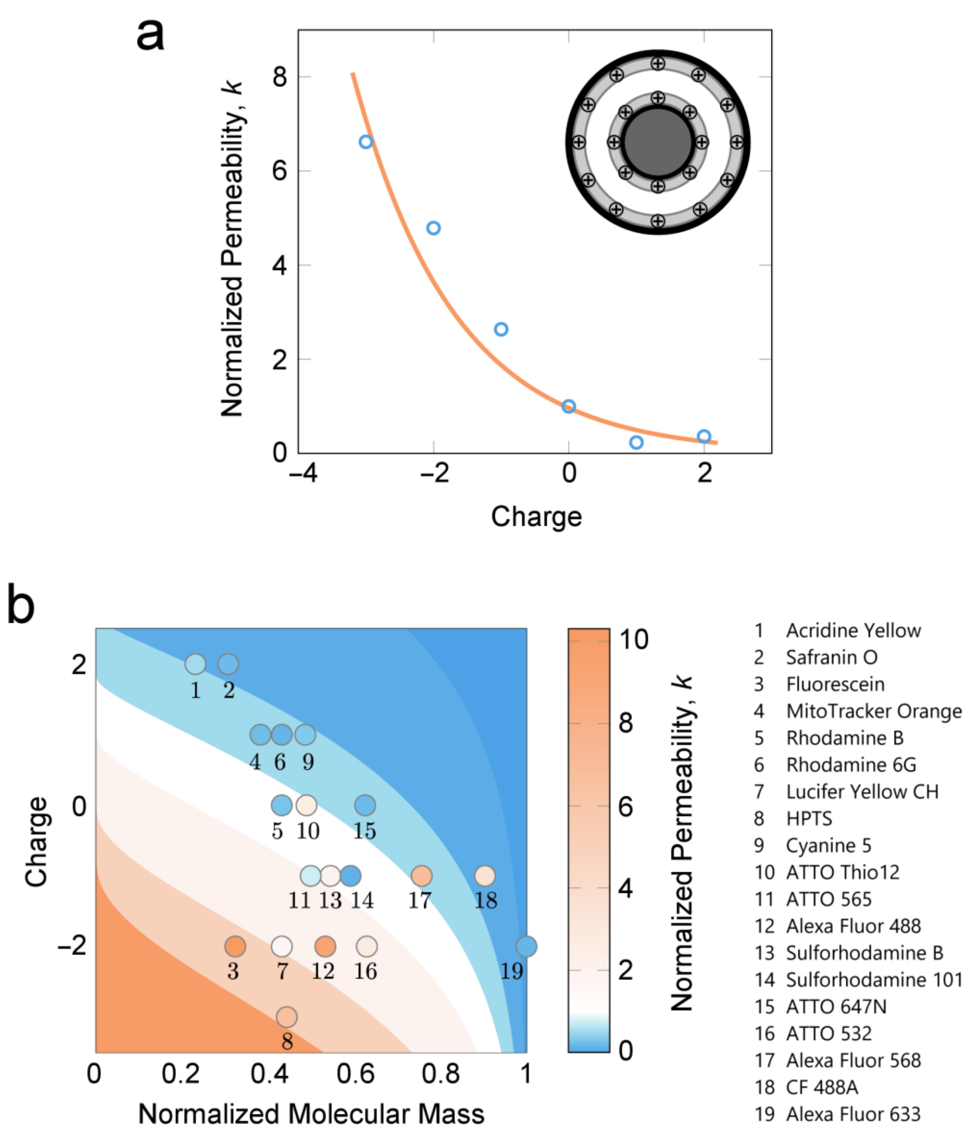


**Supplementary Figure 4. Comparison of experimentally observed plasmodesmal permeabilities, *k*, with theory. a**, Plot of fluorophore permeabilities normalised by the average permeability of the neutral fluorophores (circles) as a function of charge; values are the means of the medians of the biological replicates for each charge. The line shows a fit of the normalised theoretical permeability as a function of charge (Eqs. (3) and (5) in Ref. 1) with the surface potential ζ as the only fitting parameter. The surface potential is found as ζ = 40 mV. The theory in Ref. 1 was modified to include the particle size by changing the limits of the integral in the diffusion cross-section in Eq. (3) to the area available to the centre of a spherical particle of radius *s*. The theory is normalised by the neutral case, with the particle size found using the mean molecular weight of the neutral fluorophores. The sizes of the fluorophores are estimated from their molecular weight, assuming the largest fluorophore has a diameter equal to the cytoplasmic sleeve width $h$. We used $h$ = 3 nm and the thickness of the electrical double layer $\lambda_D$ = 1 nm (corresponding to a bulk electrolyte concentration of approximately 100 mM at room temperature; Ref. 2). The concentric geometry is approximated by a parallel plate geometry as the gap between the pore wall and the desmotubule is much smaller than the radii of either. We have simplified the surface charge distribution as shown in **a** and assume the surface potential is the same on both surfaces. The theory is valid when the concentration of the diffusing fluorophores is much smaller than the concentration of ions in the bulk cytosol, so that the presence of the fluorophores does not affect the size of the electrical double layer. This is the case when the ratio $(c_0 Z^2)(c_{el} Z_{el}^2)^{-1} \ll 1$, where $c_0$ and $Z$ are the concentration at zero surface potential and the valence of the fluorophores, respectively, and $c_{el}$ and $Z_{el}$ are the concentration at zero surface potential and the valence of the ions in the bulk cytosol, respectively (see Ref. 1 for details). Here, the ratio is approximately $(c_0 Z^2)(c_{el} Z_{el}^2)^{-1} \approx$ 0.001—0.01. **b**, Contour plot of the normalised diffusion cross-section (Eqs. (3) and (5) in Ref. 1, modified to include particle size) as a function of fluorophore charge and normalised molecular mass. The colours of the circles represent the normalised median permeabilities of the 19 fluorophores tested. The same normalisation as in panel **a** is used, along with $h$ = 3 nm, $\lambda_D$ = 1 nm, and surface potential $\zeta$ = 40 mV. The isolines are 10, 5, 2, 1, 0.5, and 0.1.

1. Christensen, A.H., Gupta, A., Chen, G., Peters, W.S., Knoblauch, M., Stone, H.A. & Jensen, K.H. Locally optimal geometry for surface-enhanced diffusion. *Phys. Rev. E* **108**, 045101 (2023).

2. Schoch, R.B., Han, J. & Renaud, P. Transport phenomena in nanofluidics. *Rev. Mod. Phys.* **80**, 839 (2008).
